# The ecological consequences of microbial metabolic strategies in fluctuating environments

**DOI:** 10.1101/2023.07.24.550395

**Authors:** Zihan Wang, Akshit Goyal, Sergei Maslov

## Abstract

Microbes adopt a variety of metabolic strategies to consume resources in fluctuating environments, but most work has focused on understanding these strategies in the context of isolated species, rather than diverse natural communities. We systematically measure the feasibility, dynamical and structural stability of multispecies microbial communities adopting different metabolic strategies. Our results reveal key distinctions between the ecological properties of different metabolic strategies, showing that communities containing sequential utilizers are more resilient to resource fluctuations, but are less feasible than co-utilizing communities.

---

Microorganisms use a variety of metabolic strategies to utilize resources in their environments. Microbial species typically either use resources all at once (co-utilization) or one after another (sequential utilization) [1, 2], with most studies focusing on the physiological benefit of using a certain strategy for an individual species [3, 4]. However, these studies tend to overlook an important ecological fact: in nature, species must coexist and compete for the same resources [2, 5]. The ecological implications of various metabolic strategies, particularly in environments where resource availability fluctuates over time, remain largely unexplored [3, 6, 7]. More specifically, we do not understand how likely communities of organisms with different metabolic strategies are to successfully and stably assemble (feasibility and dynamical stability) [8, 9, 10, 11, 12], and whether there are any trade-offs or synergies between assembly success and resilience against environmental changes (i.e., structural stability).

Here, we quantify the feasibility, dynamical stability, and structural stability of newly established microbial ecosystems, in which species implement diverse metabolic strategies, from co-utilization of resources to sequential usage. We differentiate between different sequential strategies based on how organisms adjust their physiology: “smart sequential” species utilize resources in descending order of their growth rate, while “random sequential” species use resources in a random order, disregarding their individual growth rates.

Our findings reveal that communities of smart sequential species are ∼10-fold more structurally stable than communities of co-utilizers, but have lower feasibility. That is, even though communities of smart sequential species can successfully assemble slightly less often than communities of co-utilizers, they can do so in a broader range of resource environments. Interestingly, we also found that communities containing a mix of sequential and co-utilizing species are less feasible than their “pure” counterparts, which consist only of sequentially or co-utilizing species. However, this can alleviated by giving co-utilizers a slight growth advantage. This suggests community-level competition between strategies.

We model microbial community dynamics in boom-and-bust environments. The “boom” phase is characterized by the addition of *n*_*R*_ substitutable resources at large concentrations *R*_*k*_ (much larger than their half-saturating constants), while the bust is characterized by a sudden dilution of all species by a factor *D*. This is a justifiable choice since many natural and laboratory microbial environments display boom-and-bust resource dynamics, e.g., soil [13], ocean [14] and gut microbiomes [15], and serial dilution experiments [16]. Moreover, unlike static environments (e.g., chemostats [7]), an emerging body of evidence suggests that ecological dynamics can differ significantly in such boom-and-bust environments due to the introduction of new “temporal niches” [17, 18, 19].

We model the initial stages of the colonization of a new environment by a pool of n_*S*_ species. Due to competitive exclusion, we will focus on a saturated community where the number of species n_*S*_ is equal to the number of resources n_*R*_. We assume that each species is a generalist and can consume all n_*R*_ resources, but species can utilize these resources through 3 distinct strategies: (1) co-utilizers consume all available resources simultaneously, (2) smart sequential utilizers consume resources one a time, in descending order of their growth rates, and (3) random sequential utilizers consume resources one at a time in a specific order that is uncorrelated with the growth rates on different resources.

We first quantify the feasibility — the possibility of coexistence — of “pure” communities comprising species which all adopt the same metabolic strategy. In a dynamic environment, available resources are depleted in a specific order determined by the collective resource utilization by all species [20]. This in turn influences the coexistence of those species. We determined the feasibility for a large sample of possible ecosystems, each seeded with a random set of species. For a given set of species, we asked whether there was any depletion order of resources in which they could coexist (Methods). We calculated the feasibility of pure communities with a certain metabolic strategy as the fraction of sampled ecosystems where species could stably coexist in at least one depletion order. We also developed a mathematical theory to compute the feasibility (given by 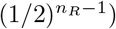 for a set of sequentially utilizing species with identical resource preferences (Methods). This theory also applies to communities under a constant resource supply (chemostats). For convenience, we measured the relative likelihood of coexistence, by normalizing the feasibility by our theoretical expectation (Fig. 1b). Since feasibility does not guarantee dynamically stability to small perturbations in species abundances, we developed a theoretical method to assess the dynamical stability of any feasible community (Methods), and only counted those communities that were both feasible and dynamically stable.

**Figure 1:**
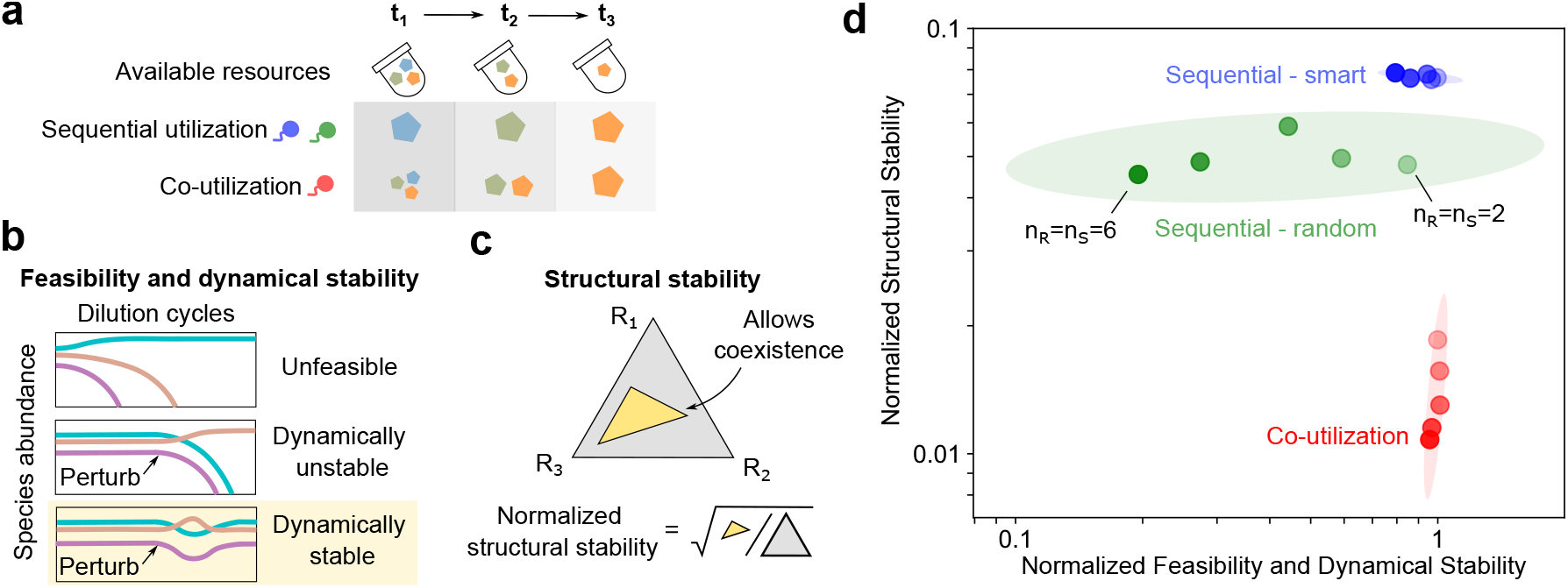
Feasibility and stability of communities adopting different metabolic strategies. (a) Illustration of sequential and co-utilizing (red) metabolic strategies. Sequential strategies can be of two types: smart (blue), where sequential resources preferences are ordered by growth rate, and random (green); (b) Schematic depicting feasibility and dynamical stability of a community; to successfully assemble, a community must be both feasible and dynamically stable; (c) Structural stability quantifies the fraction of supplied resource concentration ratios in which a feasible community can assemble; (d) Scatter plot of the feasibility and dynamical stability (normalized to theoretical expectation (Methods)) and normalized structural stability of different pure metabolic strategies. Darker colors (same as (a)) indicate a larger community complexity (number of resources and species).

While feasibility tells us whether a given set of species can in principle stably coexist, it does not ask what range of environmental conditions this coexistence will be possible in. To quantify the latter, we also developed a method to calculate the structural stability of all feasible communities (Methods). Structural stability indicates the range of resource concentration ratios in which a set of species will successfully assemble.

Remarkably, all 3 metabolic strategies differ from each other in both their feasibility, dynamical stability and structural stability (Fig. 1d, where the *x*-axis shows the combined probability of communities being both feasible and dynamically stable). The difference between the strategies becomes more pronounced as the number of resources *n*_*R*_ increases from 2 to 6. Pure communities of smart sequential species are somewhat less likely to assemble, but are most structurally stable. Random sequential species are somewhat less structurally stable, but become progressively less dynamically stable with increasing *n*_*R*_, and thus less likely to successfully assemble (Fig. S2). Finally, the feasibility of co-utilizing communities is very-well described by our mathematical theory (Supplementary Text). These communities are always dynamically stable, but their structural stability significantly suffers with increasing environmental complexity *n*_*R*_. We developed a new theory which could explain why sequential utilizers were more structural stability than co-utilizers, in terms of their variance (Methods). In a nutshell, the additional structural stability arises from many sequential utilizers never consume certain resources during a growth cycle, whereas co-utilizers always consume all available resources.

Thus far, we have looked at pure communities where all species adopt the same metabolic strategy. However, natural ecosystems typically contain a mixture of species with different metabolic strategies. We investigated the feasibility and structural stability of mixed communities, each containing a 1:1 ratio of smart sequential and co-utilizing species. Remarkably, we found that mixed communities are significantly less likely to assemble than their pure counterparts (Fig. 2a). Further, the structural stability of mixed communities was positioned between pure co-utilizing and smart sequential communities, getting progressively worse with increasing *n*_*R*_. The reduced feasibility of mixed communities is due to the fact that co-utilizing species systematically lose in direct competition with sequential utilizers, as previously suggested in the case of *n*_*R*_ = 2 [20]. Our model of metabolic strategies assumes constant proteome re-allocation to use the resources currently available in the environment (Methods), but natural microbes may deviate from this optimality assumption. This could provide co-utilizers with a distinct growth rate advantage, where the growth rate on a mixture of resources would be larger than the sum of the individual resource growth rates [21]. When we introduce a growth advantage of roughly 0.5 SD higher growth rate for co-utilizers, we found that mixed communities become much more likely to assemble, with an intermediate structural stability to their pure counterparts (Fig. 2b). As the growth rate advantage of co-utilizers increases further to 1.5 SD, the feasibility of mixed communities starts to decline again, this time due to the systematic advantage of co-utilizers (Fig. 2c).

**Figure 2:**
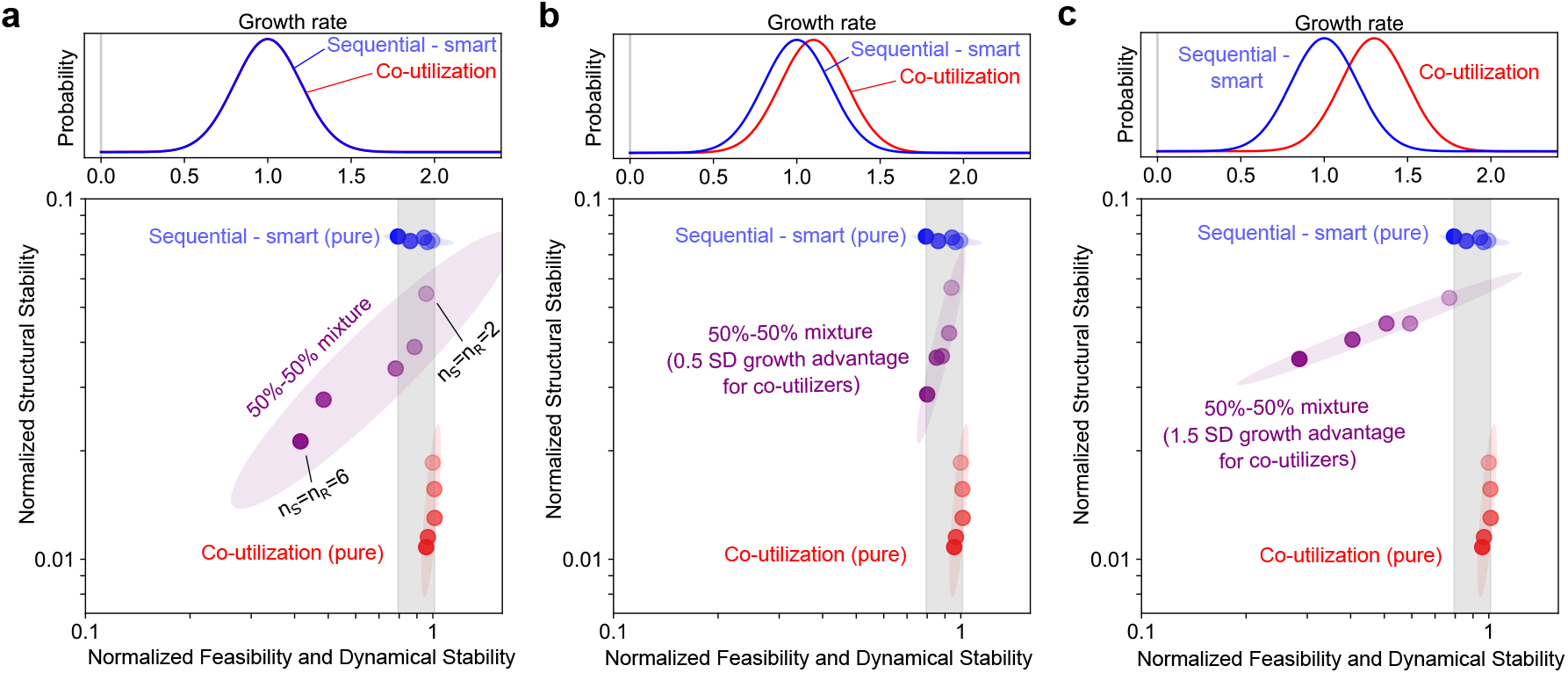
Community-level competition between sequential and co-utilizing strategies. Scatter plot of normalized feasibility and structural stability and normalized structural stability for pure and mixed (purple) communities of smart sequential (blue) and co-utilizing (red) microbes, with (a) mean co-utilizer growth rates equal to mean sequential growth rates, (b) co-utilizer mean growth rates 0.5 SD greater than sequential means, and (c) co-utilizer means 1.5 SD greater than sequential means. Feasibility and dynamical stability shows that sequential species systematically outcompete co-utilizers in (a), are about equal to each other in (b), and lose to co-utilizers in (c).

In summary, we showed that the feasibility, dynamical and structural stability of communities adopting different metabolic strategies can be drastically different. Specifically, communities of smart sequential species are the most structurally stable, but slightly less feasible than those of co-utilizers. Mixed communities containing both co-utilizing and smart sequential species are significantly less feasible, as explained by systematic growth rate differences between the two strategies.

## Acknowledgements

We thank Yulia Fridman and Yu Fu for valuable discussions and help with calculations. AG acknowledges support from the Gordon and Betty Moore Foundation under grant number GBMF4513.

## Data and code availability

There are no data associated with this paper. All code is available as a GitHub repository at the following link: https://github.com/maslov-group/ecological-consequences-metabolic-strategies

## Methods

### Model of microbial community dynamics in boom-and-bust environments

We model microbial community dynamics in a boom-and-bust environment where n_*R*_ substitutable resources are cyclically supplied at concentrations *R*_*k*_ (*k* = 1, 2, … *n*_*R*_). We will attempt to assemble a community with *n*_*S*_ = *n*_*R*_ species. Each species is assumed to be a generalist able to grow on each of these resources. We denote each species by Greek symbols, so that the abundance of species α is represented by *N*_*α*_*(t)*; the concentration of the resources k is represented by *R*_*k*_(*t*). We encode the metabolic strategy by a consumption matrix, *c*_*αk*_*(t)*, which is time-dependent. If the species α consumes the resource k at time t, we set *c*_*αk*_(*t*) to the relative fraction at which it is consumed; otherwise, we set it to 0. The consumption matrix is normalized so that at any time when the species α is growing, it has Σ_*k*_ *c*_*αk*_(*t*) = 1. Sequential utilization species consume one resource at a time, so their consumption matrices are binary and the currently consumed resource with *c*_*αk*_*(t)* = 1 is determined by their resource utilization hierarchy.

While consuming a resource, we assume that the available resource concentrations are much higher than the species’ half-saturating substrate concentrations, so that each species grows at a constant growth rate, Σ_*i*_ *g α*_*k*_*c(t)*_*αk*_ independent of resource concentrations. Thus, the dynamics of species abundance in our model can be written as follows

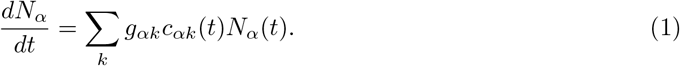

Similarly, the resource depletion dynamics can be written as follows

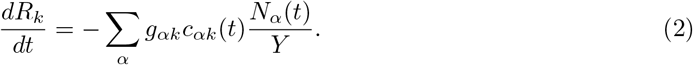

For the sake of simplicity, we will set the yield *Y* equal to 1.

The “boom” phase of the boom-and-bust cycle ends when all resources are depleted. It is then followed by a “bust” phase during which all microbial abundances are reduced (diluted) by the same factor *D* > 1 until the arrival of the next nutrient bolus, when the dynamics of the boom phase resume. Our dynamical equations are inspired by the protocol of serial dilution experiments performed in many laboratories to study microbial communities in vitro. A version of this dynamics is also realized in real-world microbial communities living in boom-and-bust environments characterized by long intervals between and identical dilution factors during the bust phase and large boluses delivered at the beginning of each boom phase.

We are interested in computing the steady state of the boom-and-bust dynamics in which each species grows by exactly the same factor *D* by which it is subsequently diluted. To compute this steady state dynamics, it is convenient to divide the boom phase of the cycle into a sequence of *n*_*R*_ *temporal niches* where only a particular subset of resources is present [20, 17, 19]. The total number of possible temporal niches for *n*_*R*_ resources is given by 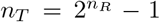. Here we have excluded a temporal niche where all resources are absent and therefore no species can grow. The resources present in the environment disappear one by one, resulting in a particular subset of *n*_*R*_ temporal niches (out of *n*_*T*_ possible ones) realized for each of the *n*_*R*_! orders of resource depletion. For example, if three resources disappear in the order 2 →3→ 1, the corresponding temporal niches are 111→ 101→ 100 (again, we do not show or count the last niche 000, which separates the end of the boom phase and the beginning of the bust phase of the cycle).

An additional complication for real species is the existence of lag times *τα* (*i, j*) when the species switches from temporal niche j to temporal niche i. Lags are not limited to the sequential resource utilization strategy, but also exist for co-utilizing metabolic strategies (see e.g. one of the two species studied in ref. [22]). During the lag time, the species *α* does not consume any resources, so Σ_*i*_ *c*_*αi*_(*t*) = 0. For simplicity, in most of the mathematical calculations shown below we will set the lag time *τ*_*α*_ = 0 and comment on a more general case only when necessary.

was

### Details of numerical simulations

In our numerical simulations we used *D* = 100, and *gα*_*i*_ = *g*_*0*_ +*σ*_*g*_*Zα*_*i*_, where *Zα*_*i*_ is a random number selected form the standard normal distribution. For figures used in the main text of the study we used *g*_0_ = 1 hr^*−*1^ and *σ*_*g*_ = 0.2 hr^*−*1^. For these parameters one does not need to take special precautions that the growth rate remains positive since negative rate would require *Zα*_*i*_ < −5, which is a very unlikely. In supplementary figures where we used larger values of *σ*_*g*_. To keep our growth rate distribution symmetric around its mean we symmetrically truncated the normal distribution so that 0 < *gα*_*i*_ < 2. Simulations were carried out in Python (see Code Availability Statement). Every metabolic strategy was sampled 10,000 times.

### The feasibility of a community assembly

The steady state of a microbial community in a boom-and-bust environment is characterized by a particular depletion order of resources which depends on abundances of both species and resources at the start of the boom cycle in a complex fashion (see later sections for details). Since it is not known a priori in which order the resources might be depleted, one should test all *n*_*R*_! possible depletion orders for feasibility. For a given resource depletion order let *G*_*αi*_ be the growth rate of the species α in the temporal niche *i* ranging from *i = 1* where all *n*_*R*_ resources are present to *i = n*_*R*_ where only one resource remains. Let *t*_*i*_ be the duration of each of these temporal niches. In the absence of time lags we can calculate the growth ratio of each species during the entire boom phase of the cycle as exp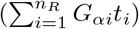. In the steady state, this ratio must be equal to the dilution factor *D* during the bust phase of the cycle, which leads to the following equations for each of *n*_*S*_ species

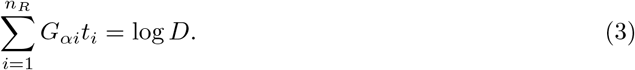

The matrix *Gα*_*i*_ used in this equation is related to the matrix *gα*_*k*_ of growth rates on individual resources, but this relationship depends on the metabolic strategy as described in the next section.If the matrix *Gα*_*i*_ is invertible (that is, the duration of each temporal niche is determined by

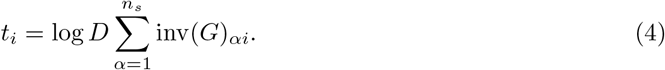

This solution is biologically and physically feasible provided that all time *t*_*i*_ > 0. In principle, after checking all *n*_*R*_! depletion orders for feasibility one can get more than one feasible solution, corresponding to different ways of assembling the microbial community on a given set of resources. In practice, in our simulations of sequential metabolic strategies with *n*_*S*_ *= n*_*R*_ = 4 we never observed more than 5 feasible depletion orders for the same community. Even these often turned out to be dynamically unstable (for definition of dynamical stability, see the corresponding section). Feasibility of assembly of a given community has a useful geometrical interpretation. Consider one specific nutrient depletion order. We can rescale the columns of the matrix *G α*_*i*_ by subtracting from each column the sample mean of growth rates 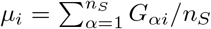 of all species in this niche and dividing the resulting value by *n*_*S*_*σ*_*i*_. Here the standard deviation *σ*_*i*_ of the sample is defined by 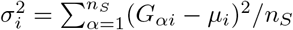

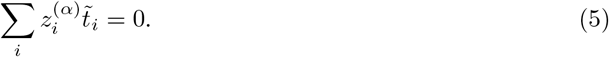

Here 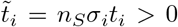. Note that after this rescaling, the vectors 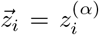 have unit length and lie on the hyperplane defined by 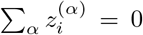. Hence they all lie on the (*n*_*S*_ − 1)-dimensional unit hypersphere centered at the origin given by the intersection between the *n*_*S*_-dimensional unit hypersphere centered at the origin and this hyperplane. The assembly of a given set of species is feasible in a given depletion order if and only if the origin (0, 0, …, 0) lies inside the convex hull of *n*_*R*_ vectors 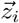.

### Metabolic strategies

A species using sequential nutrient utilization strategy is characterized by a hierarchical ranking of resources *h*_*αk*_ ranging from *h*_*αk*_ = 1 (meaning *R*_*k*_ is the top choice of the species α) to *h*_*αm*_ *= n*_*R*_ (meaning that *R*_*m*_ is the the bottom choice of the species α). In each temporal niche, a sequentially utilizing species will exclusively consume the highest ranked resource (one with the lowest *h*_*αk*_ rank) among all resources present in this niche. In this study, we consider two different sequential utilization strategies:

- “Smart” sequential utilization strategy for which the hierarchy of resource utilization exactly matches the hierarchy of growth rates g_*αk*_ ranked from the fastest (rank 1) to the slowest (rank *n*_*R*_).
- Random sequential utilization strategy for which the rank of a resource is uncorrelated with the growth rate of the species on this resource.

In addition to sequential utilization strategies, we also consider species that simultaneously utilize all currently available resources. Such co-utilizing species are characterized by the vector of consumption ratios *c*_*αk*_ > 0, normalized so that Σ_*k*_ *c*_*αk*_ = 1. If all *n*_*R*_ nutrients are present in the medium (the temporal niche 11 … 1), the growth rate of a co-utilizing species is given by *Gα* all resources 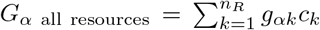. In a temporal niche where only a subset of resources remain, we assume that the organism reallocates its proteome to consume only the available resources, while keeping relative consumption ratios unchanged. In this case, the growth rate is given by 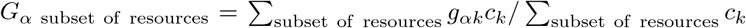. Regarding the choice of the vector of consumption ratios, we consider two metabolic strategies

- Uniform co-utilization strategy with *c*_*αk*_ = 1/*n*_*R*_.
- Random co-utilization strategy with species-dependent *c*_*αk*_ given by uniformly distributed random numbers normalized by their sum.

In this study we present the results for the uniform co-utilization strategy, while the results for a random co-utilization strategy are very similar. Note that in the case of an extremely broad distribution of c_*k*_’s, the co-utilization strategy turns into the sequential utilization strategy with preferences determined by the ranking of c_*k*_. Indeed, if one of the c_*k*_ is much larger than others, the microbe will almost exclusively use that resource until it runs out, at which point the proteome will be reallocated and the resource with the second largest c_*k*_ will take its place. By choosing *c*_*αk*_ = exp(λ*g*_*αk*_)/ Σ_*m*_ exp(λ*c*_*αm*_) one can study a gradual transition between the uniform coutilization strategy for λ = 0 and the smart sequential utilization strategy for λ ≫ 1.

### Theoretical expectation of the feasible assembly success rate

The mathematical theory [23, 24] that allows one to calculate the probability of a feasible assembly for a randomly selected set of species and nutrients has been developed only if the vectors 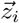 (normalized columns of *G*_*αi*_) are independently chosen from a distribution that is symmetric with respect to a transformation 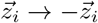. This is not true for the general case of microbial communities assembled in boom-and-bust environments, where columns of the matrix *G*_*αi*_ are correlated in a non-trivial way. For example, the growth rate of sequentially utilizing microbes may remain the same over multiple temporal niches as long as the currently used resource does not disappear from the environment. If the first choice of a microbe is the last resource to be depleted, *G*_*αi*_ will remain the same in all *n*_*R*_ niches.

However, in a certain special case, where random sequentially utilizing species all have identical resource preference order, which we can assume without loss of generality to be *h*_*αi*_ = *i*. In this case, the only possible order of resource depletion is 1 → 2 → … n_*R*_, and the matrices *G*_*αi*_ and *g*_*αi*_ are identical. Indeed, in the first temporal niche, each species α consumes the resource *R*_1_ at the growth rate *g*_*α1*_, in the second temporal niche - the resource *R*_2_ at the growth rate *g*_*α1*_, and so on. If the columns of the matrix *g*_*α*1_ are not correlated with each other, and if they are selected from some distribution that is symmetric about its mean (possibly different for different resources), then the conditions species in Ref. [23] are satisfied. In this case, the probability *P*_*feasible*_ of finding a feasible solution in a more general case where *n*_*R*_ - the number of resources and hence the number of vectors 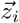 lying on the (*n*_*S*_ − 1)-dimensional unit sphere - may be different from the number of species is given by [23]

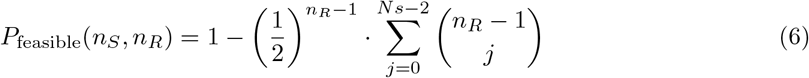

In one case considered in this article, 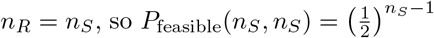

The assumption used in deriving eq. (6) is that for every resource i the distribution of species growth rates *g*_*αi*_ is symmetric around its mean value. Any deviation from the symmetry 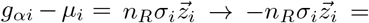, such as e.g. in the case of sparse matrices *g*_*αi*_, has been shown [24] to decrease the probability of finding a feasible solution. Another assumption is that the elements of the consumption matrix for different resources are chosen independently from each other. This means, for example, that there are no trade-offs between *G*_*αi*_ for different resources i that would make feasible solutions more likely. In one special case of a perfect linear tradeoff Σ*i G*_*αi*_ = const the probability of a feasible assembly becomes equal to 1. Indeed, similar to the chemostat community studied in Ref. [25], the boom-and-bust community composed of an arbitrary number of species with a perfect tradeoff can in principle coexist with each other in a state with equal temporal niche duration *t*_*1*_ *= t*_2_ = … *t*_*nR*_. Conversely, positive correlations between growth rates of the same species on different resources would make feasible solutions less likely. They can be implemented in our model by introducing a species-specific mean value µ_*α*_ for growth rates of a given species on each of the resources.

### Structural stability with respect to resources

A feasible community has a unique set of temporal niche durations *t*_*i*_ > 0. The next step is to find the range of relative resource concentrations *R*_*i*_ in the nutrient bolus that lead to community assembly. This can be done using the mass conservation rules in the resource-to-biomass conversion. The temporal niche time intervals *t*_*i*_ fully determine the factors *F*_*αi*_ = exp(*G*_*αi*_*t*_*i*_) by which individual microbial species grow during that temporal niche. During this time, the biomass of the species α increased from П_*j<i*_ *F*_*αj*_ to *F*_*αi j*_П_*<*i_ *F*_*α*j_. This increase in biomass is equal to the total resources consumed during this temporal niche. The amount of resource *R*_*k*_ consumed is given by *N*_*α*_(0)*c*_*αk*_(during niche i)(*F*_*αi*_ − 1) П_*j<i*_ *F*_*αj*_. This allows one to compute the *M*_*αk*_ matrix, which converts species abundances *N*_*α*_(0) at the start of the boom phase into resource quantities they consumed during the entire boom phase.

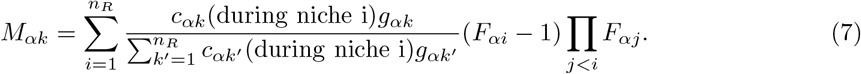

Due to mass conservation column sums are given by 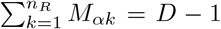. Indeed, in the steady state, the abundance of each species must increase by a factor *D*. This extra biomass given by *N*_*α*_(0)(*D* − 1) must be equal to the total quantity 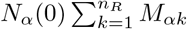 of all resources consumed by this species.

The structural stability *S*_*R*_ of a feasible community can be quantified by the fraction of all bolus resource ratios for which the community successfully assembles. It is proportional to the determinant | det(*M*)|. To properly normalize it, one must first divide the matrix *M* by *D* − 1 so that the sum of the elements in each column is equal to 1. If one randomly chooses *n*_*R*_ − 1 ratios between resource concentrations *R*_*k*_ in the nutrient bolus at the beginning of each boom phase of the cycle, the fraction of the volume of the simplex Σ_*k*_ *R*_*k*_ = 1, *R*_*k*_ > 0 that results in community assembly is given by

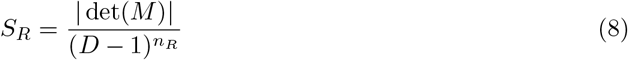

The structural stability defined in this way exponentially scales with the number n_R_ −1 of independent ratios between resource concentrations. The normalized stability defined by

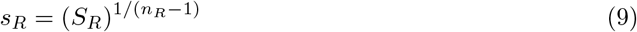

corrects for this effect. As can be seen from this study, for *D* = 100, g_0_ = 1 and *σ*_*g*_ = 0.2 used in our study, the average value of *s*_*str* R_) in communities composed of sequentially utilizing species (both smart and random) is approximately independent of *n*_*R*_. The intuitive interpretation of this quantity is the approximate range of individual nutrient ratios that result in successful community assembly.

In a general case normalized structural stability depends on *n*_*R*_ (the number of resources in the environment), the dilution ratio *D*, the average growth rate *g*_0_ and the standard deviation σ_*g*_ of growth rates in the *g*_*αi*_ matrix (see Fig. S1).

### Dynamical stability

To determine the dynamical stability of a given community, we examine how a small perturbation in the abundance of species α at the start of one boom-and-bust cycle, *N*_*α*_(0), affects abundances of all species β at the start of the next cycle, which we denote by *N*_*β*_(0 at the next cycle). We construct and analyze a sensitivity Jacobian matrix *J*_*αβ*_ defined by *J*_*αβ*_ = ∂*N*_*β*_(0 at the next cycle)/∂*N*_*α*_(0). The community is dynamically stable if the real parts of all eigenvalues of *J*_*αβ*_ have absolute values less than 1, and dynamically unstable otherwise.

The elements of *J*_*αβ*_ can be computed in the following way:

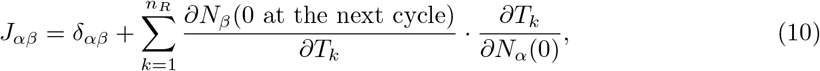

where *T*_*k*_ is the time from the start of the cycle until resource *R*_*k*_ s depleted. Note that it is different from the duration of the temporal niche *j(k*) during which this resource is depleted. Let *k(j)* to mark the resource that was depleted at the end of the temporal niche *j* and *j(k)* - the niche at which the resource *k* was depleted. For example, if the depletion order of resources is *R*_3_ →*R*_1_ →*R*_2_, then *k*(1) = 3, *k*(2) = 1, and *k*(3) = 2. Conversely, *j*(1) = 2, *j*(2) = 3, and *j*(3) = 1. For convenience of notation we will also define *k*(0) = 0 and *T*_0_ = 0.

The abundance of species *β* at the start of the next cycle, *N*_*β*_(0 at the next cycle), can be written as:

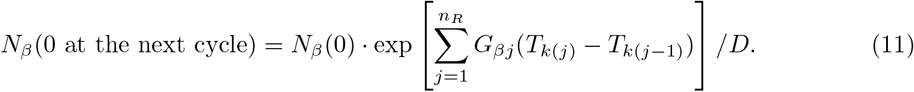

Indeed, in the new notation the duration *t*_*j*_ of the temporal niche *j* can be written as the difference between *T*_*k*(*j*)_ (the time until depletion of the nutrient that disappeared during niche j) and *T*_*k(j−1)*_ (the time until depletion of another nutrient that disappeared during the previous niche *j* − 1). We will use Eq. (11) to calculate *∂N*_*β*_(0 at the next cycle)/∂*T*_*k*_. Only two temporal niches: the niche during which the resource *R*_*k*_ was depleted and the niche immediately following it contribute to this partial derivative. The growth rates of a species *β* in these two niches is given by *G*_*β,i*(*k*)_ and *G*_*β,i*(*k*)+1_, respectively. Note that in the steady state one has exp 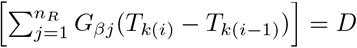. From this one gets

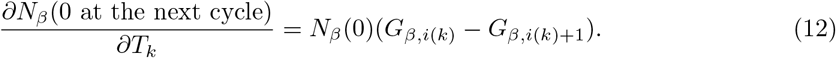

Here 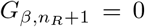 is the growth rate in the final temporal niche where all nutrients have been depleted.

To calculate ∂*T*_*k*_/∂*N*_*α*_(0) we use the matrix *M*_*αk*_ converting the species abundance at the start of the cycle to resource concentration *R*_*k*_ in the steady state

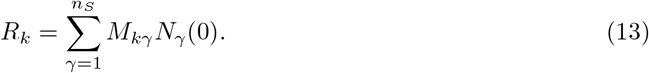

Resource concentrations *R*_*k*_ supplied at the start of every cycle are held constant and thus independent of species abundances. Hence, a partial derivative of the left hand side of Eq. (13) with respect to *N*_*α*_(0) is equal to 0. Making the partial derivative of the right hand side of Eq. (13) equal to 0 results in the following equation

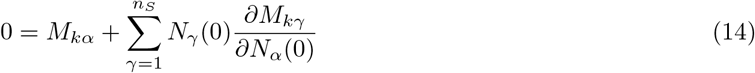

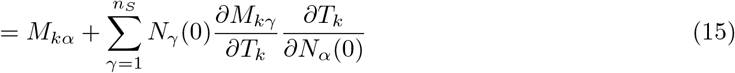

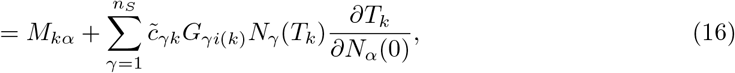

Here we used the fact that *M*_*γk*_ depends on *T*_*k*_ only for species *γ* that were using *R*_*k*_ when it was depleted. The contribution of these species to *N*_*γ*_(0)*M*_*γk*_ is equal to their abundance *N*_*γ*_(*T*_*k*_) times the consumption fraction 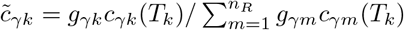. Because *N*_*γ*_(*T*_*k*_) scales with *T*_*k*_ as exp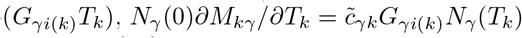.

From Eq. (16) one gets

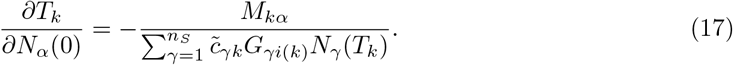

By plugging Eq. (12) and Eq. (17) into Eq. (10) one finally gets

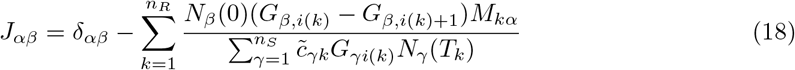

Note that the Jacobian matrix depends not only on the order of nutrient depletion, but also on the steady-state species abundances. Thus, in principle, a given feasible solution may be dynamically stable for some species abundances, but not for others.

The condition abs(Re(λ)) < 1 guarantees the dynamical stability of the steady state for small fluctuations in species abundances. However, this analysis does not prevent the microbial community from transitioning to a multi-cycle or even chaotic dynamical state, as described in Ref. [19]. In agreement with the results of Ref. [19], we observe that in the absence of fluctuations in resource concentrations such behavior is very rare (less than 1 in 10^4^ runs) in our simulations of *n*_*S*_ = *n*_*R*_ = 4 and therefore makes only a negligible correction to our main results.

## Supplementary Figures

**Figure S1:**
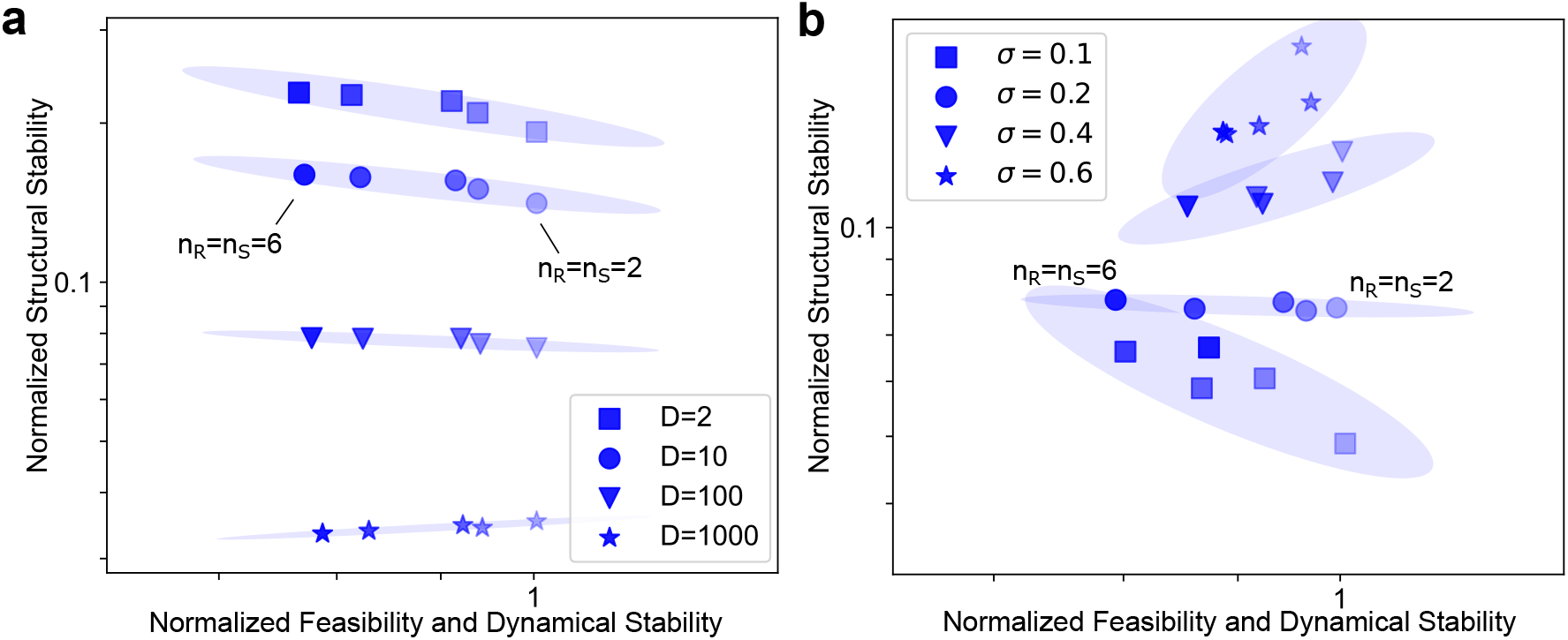
Dependence of ecological properties on the dilution factor D and the standard deviation *σ*_*g*_ of growth rates. Scatter plots of the normalized feasibility and dynamical stability and the normalized structural stability of smart sequentially utilizing species as a function of different parameters. Each point reflects the average over 10,000 randomly generated communities. Darker colors of the same shape imply higher *n*_*R*_ and *n*_*S*_, and shapes represent different values of the corresponding parameter (as in legend). (a) Dependence of feasibility, dynamical and structural stability on (a) the dilution factor D, and (b) the standard deviation of the growth rate distribution *σ*_*g*_ (Methods).

**Figure S2:**
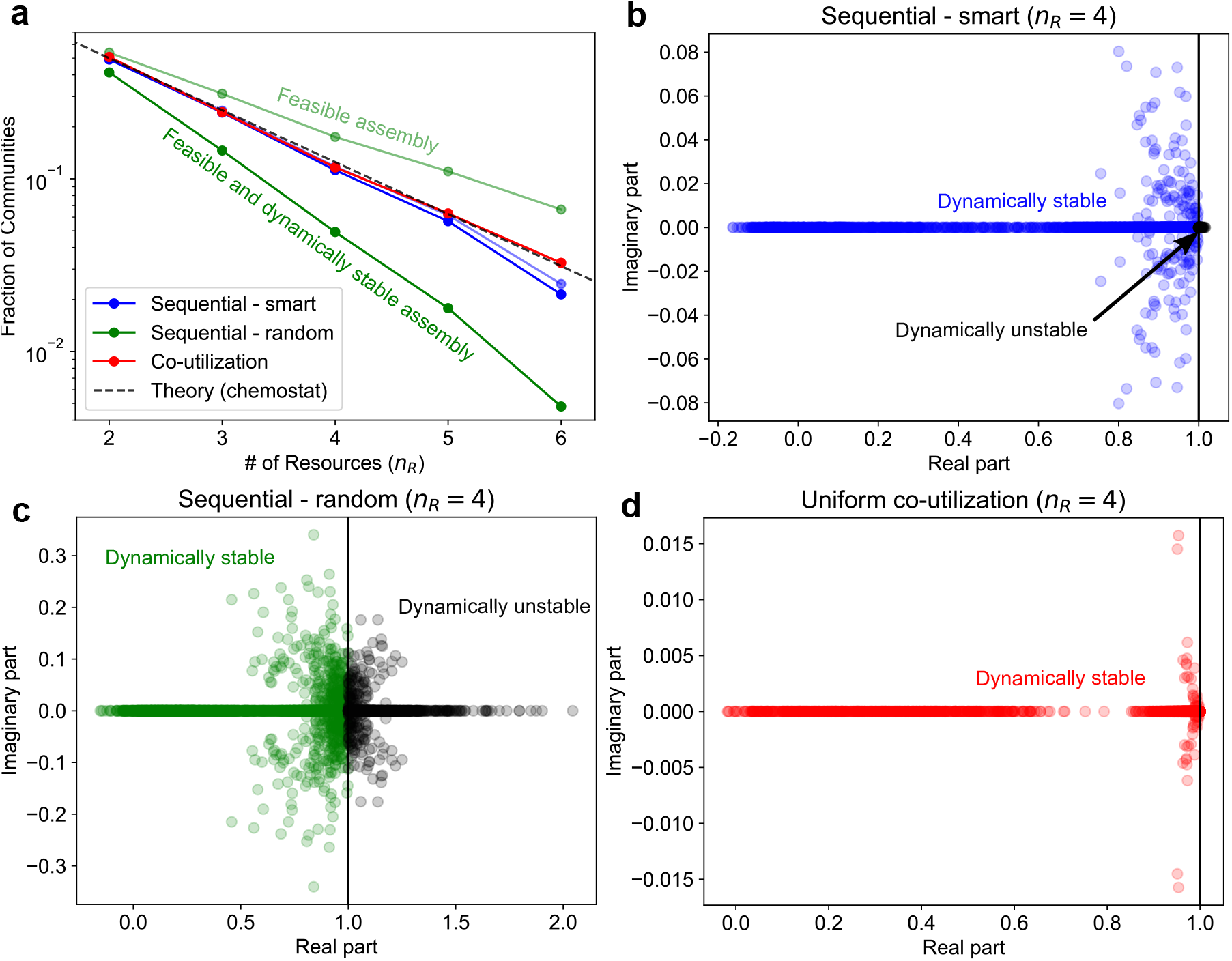
Dynamical stability of different metabolic strategies. (a) Plot of the fraction of feasible and dynamically stable communities seen in simulations with the number of resources *n*_*R*_ (*n*_*R*_ *= n*_*S*_). Each point indicates results from 10,000 randomly generated communities. Different colors represent different metabolic strategies (see legend); lighter colors indicate all feasible communities, irrespective of dynamical stability (see Methods eqn. 4); darker color indicates both feasible and dynamically stable communities. The dashed black line indicates the theoretical expectation, i.e., 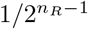 (Methods). (b)-(d) Eigenvalues of the sensitivity Jacobian matrix J (Methods) for different metabolic strategies, for all feasible communities in (a). Feasible but dynamically unstable communities (black dots) are (b) rare for smart sequential species, (c) common for random sequential species, and (d) absent for co-utilizers.

